# The demand for wildlife not protected by the CITES multilateral treaty

**DOI:** 10.1101/2022.03.03.482781

**Authors:** Freyja Watters, Oliver Stringham, Chris Shepherd, Phillip Cassey

## Abstract

The international wildlife trade presents severe conservation and environmental security risks. However, no international regulatory framework exists to monitor the trade of species not listed in the appendices of the Convention on International Trade in Endangered Species of Wild Fauna and Flora (CITES). We explored the composition and dynamics of internationally regulated versus non-regulated trade, focussing on importations of wild-caught terrestrial vertebrates entering the United States of America (US) from 2009-2018. The prominence of the US in global wildlife imports and its detailed data collection conventions allows a unique opportunity to formally assess this substantial but often overlooked and understudied component of the legal wildlife trade. We found 3.6 times the number of unlisted species in US imports compared with CITES-listed species (1,366 versus 378). CITES-listed species were more likely to face reported conservation threats relative to the unlisted species (71.7% vs 27.5%). Yet, we found 376 unlisted species facing conversation threats, 297 species with unknown population trends and 139 species without an evaluation by the IUCN Red List of Threatened Species. Unlisted species appeared novelly in imports at 5.5 times higher rates relative to CITES-listed species, where unlisted reptiles saw the largest rate of entry, averaging 53 unique species appearing in imports for the first time per year. Overall trade volumes were substantially larger for unlisted imports with approximately 11 times the number of animals relative to CITES-listed imports, however, import volumes were similar when compared at a species-by-species level. We found that the countries that were top exporters for CITES-listed shipments were mostly different from exporters of unlisted species. In highlighting the vulnerabilities of the wild-caught unlisted vertebrate trade entering the US and in the face of increasing global demand, we recommend governments adapt policies to monitor the trade of all wildlife.

## Introduction

The vulnerability of large-bodied charismatic species to overharvesting and exploitation from the wildlife trade has attracted considerable scientific and popular attention (Cardoso et al., 2021). Yet, the risk that the international wildlife market poses to the survival of many lesser-known or less charismatic species is frequently overlooked (Fukushima et al., 2020; Margulies et al., 2019). Annually, tens of thousands of species are traded globally to supply the widespread demand for traditional medicines, food, luxury items, exotic pets, and plant cultivation (Harfoot et al., 2018). Accordingly, wildlife trade has emerged as one of the leading threats to global biodiversity and environmental security (Gore et al., 2019; Maxwell et al., 2016). Wild populations are particularly at risk of extinction from overexploitation when harvesting is unsustainable (Harris et al., 2016; Lenzen et al., 2012; Scheffers et al., 2019). Around 18% of all terrestrial vertebrate species have been recorded in global trade (c. 5.6k species), with traded species found to endure higher levels of extinction threat in comparison to non-traded species (Morton et al., 2021; Scheffers et al., 2019). Additionally, the transnational wildlife trade poses substantial disease risks for both wildlife and humans, a fact that has been frequently highlighted by the ongoing COVID-19 pandemic (Aguirre et al., 2020; O’Hanlon et al., 2018).

Species at risk of over-exploitation from international trade aren’t automatically designated to be protected. CITES is the largest body regulating the international trade in wildlife (CITES, 2020). Currently, of all extant described species, only 10.5% of amphibians, birds, mammals and reptiles are listed in CITES (c. 3691 species). For all species without CITES protection, no such regulatory framework exists to monitor their international trade. It is only after the documentation of major declines in wild populations, or large volumes of illegal trade seizures, that many species are identified as at risk from trade (e.g., (Bergin et al., 2017; Challender et al., 2014; Shepherd & Ibarrondo, 2005; Waeber et al., 2019). However, for many species, data on key life history and population metrics are either unavailable or incomplete, making the assessment of the anticipated risks associated with trade potentially ineffectual (Smith et al., 2011). Regulation of the trade in threatened species by CITES lags markedly behind the assessments published by the International Union for Conservation of Nature (IUCN) Red List of Threatened Species (hereafter the IUCN Red List), with over 28% of threatened Red Listed species recognized as active in international trade not currently listed in the CITES appendices (Frank et al., 2019).

Individual governments may maintain import and export records for species not listed by CITES, but the scope and availability of data are largely dependent upon the efforts and priorities of local authorities. Furthermore, few countries have historically kept any records, let alone publicly available records, of importation and exportation data for species not listed by CITES. To better understand the scale of trade (e.g., volume and diversity of species), data on the trade of unlisted species needs to be collected, and standardized. The United States of America (US) is one country that keeps a detailed database of both CITES-listed and unlisted traded species. The US Law Enforcement Management Information System (LEMIS) database, maintained by the US Fish and Wildlife Service (USFWS) retains records of all declared imported and exported wildlife of both CITES-listed and unlisted species entering and leaving the country. In terms of monetary value, the US is the largest trader of wildlife and wildlife products worldwide (Andersson et al., 2021). The data from LEMIS, therefore, offers a valuable opportunity to investigate the scope of the trade in species not protected by the CITES treaty.

Currently, no large-scale (i.e., country-level) comparison of internationally regulated trade versus non-regulated trade exists (Olsen et al., 2019). Here, we focused on importations of wild-caught amphibians, birds, mammals, and reptiles entering the US for the ten years encompassing 2009-2018. We explored the composition and dynamics of the trade in species not listed in the CITES appendices and compared trade dynamics with the trade in CITES-listed species. Specifically, we (i) assessed how trade might impact wild populations by examining species vulnerability according to IUCN Red List classifications, highlighting those threatened by extinction, those with declining populations and those threatened by intentional harvesting; (ii) compared the species composition of imports and the dynamics of species traded through time, (iii) compared the trade volumes of imports, and (iv) compared the key exporting countries to the US, highlighting those with high volumes and/or species richness of exports. We discuss the differences in trade dynamics of unregulated species as an opportunity for reform in the ways that international wildlife trade is monitored and regulated.

## Methods

### Data compilation

We obtained records for the years 2009-2018 from the USFWS LEMIS database (obtained via requests to the US government; see (Romagosa et al., 2009) for more detailed information on the database). The records provided data on imports, exports and re‐exports between the US and other countries, with each row of data representing a record for specimens or products of one taxon (i.e., species) with the same importer, exporter, shipping dates and trade term codes. In this study, we focused on records of imports of live animals entering the US from wild-caught sources. Transactions whose reported purpose was classified as ‘Commercial’ or ‘Personal’ were included in our analysis whilst trade intended for scientific, research or educational purposes were excluded. We excluded records with no identifiable species name (e.g., genus or family level identification) and excluded records with no specified country of export origin from our analysis. We standardized species names to the most recent versions of the following taxonomic databases: AmphibiaWeb (AmphibiaWeb, 2021), International Ornithological Congress (IOC) World Bird List (Gill & Donsker, 2021), Mammal Diversity Database (Mammal Diversity Database, 2021) and the Reptile Database (Uetz, 2021).

To assess if traded species were more likely to be assessed as vulnerable to extinction, we collected data on several assessment categories for species classified by the IUCN Red List (as of November 2021; (IUCN, 2021)). First, we defined species as being threatened with extinction if their Red List category was vulnerable (VU), endangered (EN) or critically endangered (CR) (IUCN 2021). Next, we recorded whether the IUCN threats classification scheme (Version 3.2) listed species as being under an ongoing threat from intentional harvest/use (code 5.1.1 intentional use: hunting & collecting terrestrial animals). We also recorded whether the species was listed by the IUCN as present in international level trade and intentional use. We described species as having a declining population if their population trend status was listed as decreasing. In addition, we determined if each species’ IUCN Red List assessment was out of date (i.e., needing updating) by recording the year of the most recent assessment. We considered assessments as out of date if they occurred greater than 10 years prior (i.e., before 2011); following the criteria used by the IUCN (IUCN 2021).

### Statistical analyses

We determined if there were differences in the proportion of CITES-listed species versus unlisted species, for each of the following IUCN assessment categories: i) red list-status (threatened/not-threatened/not-evaluated); ii) population trend (declining/stable or increasing/ unknown); and iii) threatened by intentional use (threatened/not-threatened). This was repeated for each taxonomic class. To perform these comparisons, we used contingency-type analyses, testing for statistical independence with Fisher’s exact tests (Appendix S1).

To assess whether the number of unique species imported each year for CITES-listed and unlisted trade varied significantly over the ten years the dataset spanned, we used generalized linear models (GLMs) with a Poisson distribution and with the number of unique species (per year) as the response variable and year as the predictor variable. We excluded the year 2009 from the GLM as this was used as a reference year to determine the baseline composition of species. We further explored the temporal trends of species occurrence in imports with species accumulation curves across time, which plots the number of unique species appearing in CITES-listed and unlisted imports for the first time between 2010-2018, with 2009 as the reference year.

We quantified import volumes per species by tallying the total number of imported live animals for each species over the ten-year sampling period. To evaluate whether significant differences in the total import volumes across species occurred between CITES-listed and unlisted trade, we used GLMs with the total volume of each species as the response variable and CITES status as the explanatory variable for: a) all species; and b) all species by IUCN assessment category. We tested various model distributions (Poisson, negative-binomial, Gaussian) with Akaike’s Information Criterion (AIC) to assess which would best fit the data and found the best performing model with the lowest AIC was a gaussian distribution with the response variable volume on the log^10^ scale (Appendix S7, S11). While transforming ecological count data is not appropriate in some circumstances (O’Hara & Kotze, 2010), our data did not fit these assumptions for models with either a) all species or b) all species by IUCN assessment category (i.e., no zero observations and the log-transformed gaussian model performed well due to the underlying distribution of data being extremely dispersed).

We identified the key exporting countries of origin for CITES-listed and unlisted trade over the ten-year sampling period based on: i) the total volume of trade from countries whose exporting volume to the US were ≥ 5% of total exports for that trade type and visualized these country-type relationships with Sankey diagrams, and ii) the species richness of exports with choropleth maps.

The VCD package was used for mosaic plots (Meyer et al., 2006, 2021). All plots and choropleth maps were made with the Ggplot2 package (Wickham, 2016). Sankey diagrams were made with Ggplot2 and Ggforce packages (Pedersen, 2021). All other analysis was performed with base functions in the R statistical software version 4.1.2 (R Core Team, 2021).

## Results

Over three times the number of unlisted species (1356 wild-caught species) were imported to the US compared with 378 CITES-listed species. For each taxonomic group, the number of imported unlisted species was greater than the number of CITES-listed imported species, ranging from approximately triple (reptiles, n = 803 unlisted species) to seven times (amphibians, n = 232 unlisted species) (Fig. 1).

**Figure 1.**
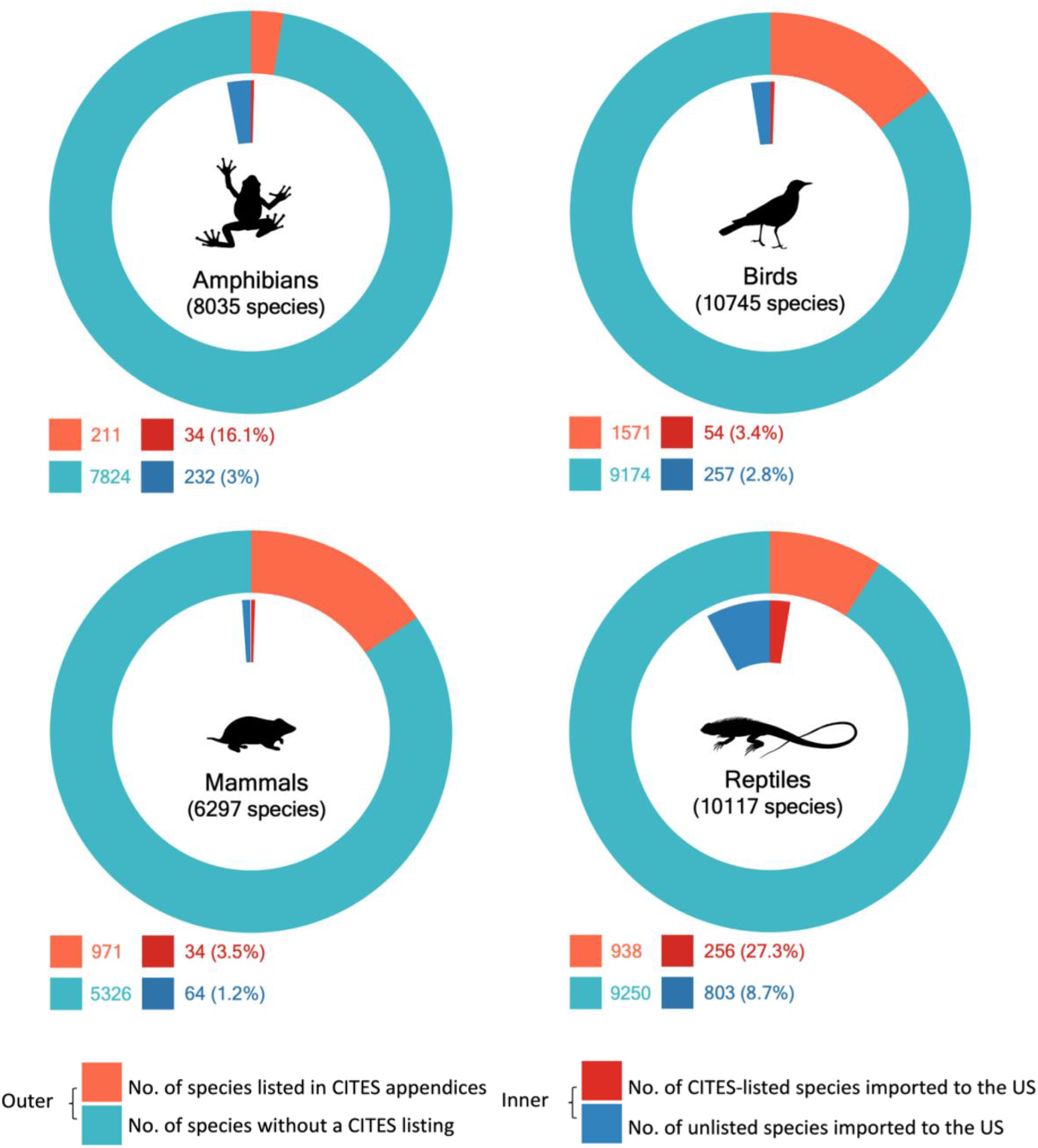
Trade in all wild-caught live CITES-listed and unlisted species in comparison to the total number of currently described extant species. The outer circles represent the total number of described species for each taxonomic class (i.e., global diversity as of 2020), divided into those that are CITES-listed and those that are not. The inner circles represent the number of listed/unlisted species that were imported to the US between 2009-2018 (from the LEMIS database) according to their current CITES status (as of December 2021).

Compared with unlisted species, CITES-listed species of all taxonomic classes were significantly more likely to be categorized by the IUCN Red List as threatened with extinction, having declining populations, and threatened by intentional use (except mammals for the latter category; Fig. 2, Appendix S1). At the same time, 376 unlisted species were categorized by the IUCN Red List as being either threatened, having a declining population, or being threatened by intentional use; with unlisted reptiles having the most species (153) followed by unlisted amphibians (96), birds (90), and mammals (37) (Appendix S2). 297 IUCN Red List listed imported unlisted species showed unknown population trends (Appendix S2) and 139 did not have an IUCN Red List evaluation (Fig. 2). Around half of unlisted species imported to the US were not recorded by the IUCN Red List as being intentionally used and also not present in international trade (Appendix S3).

**Figure 2.**
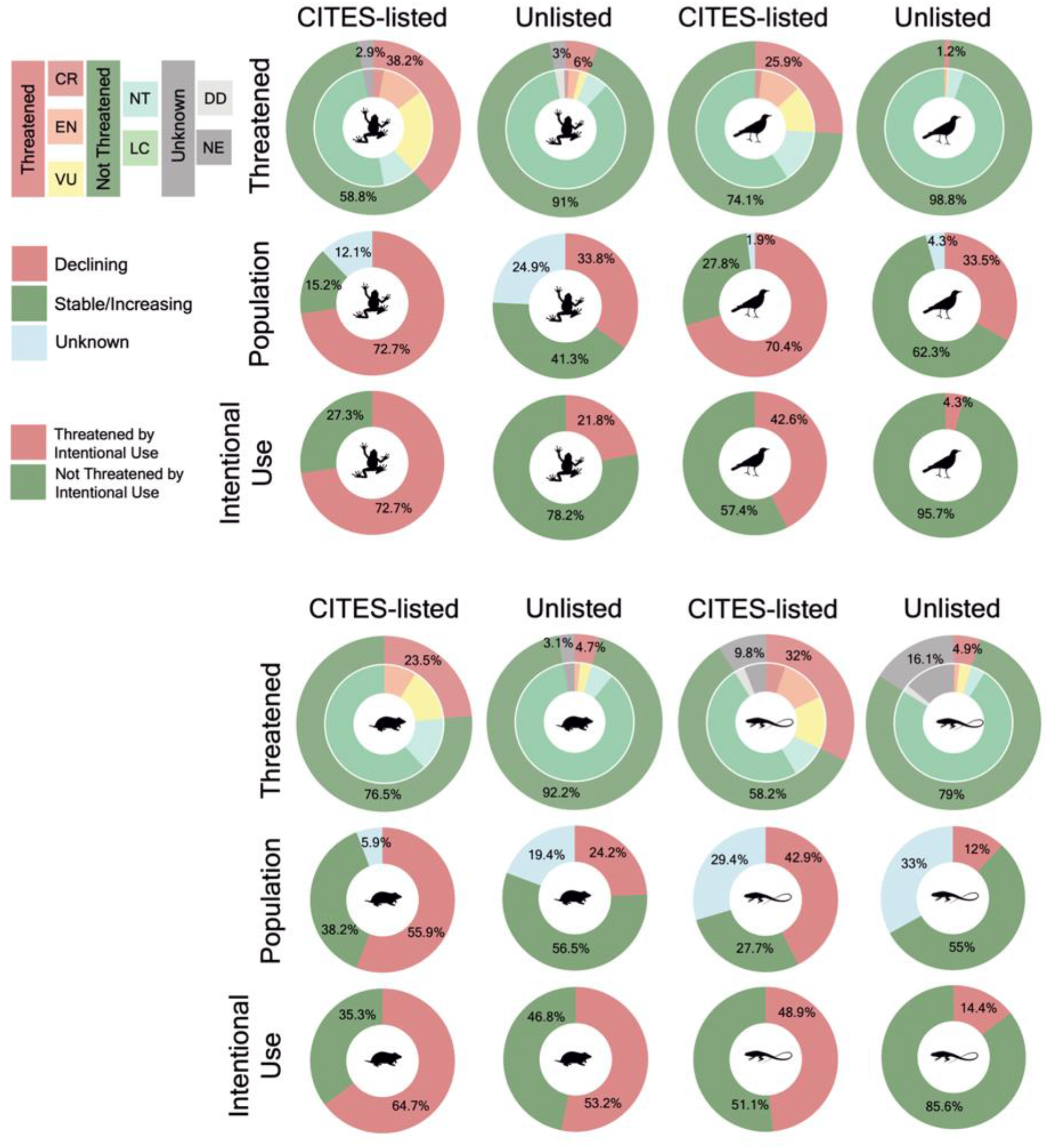
IUCN assessment category comparison between CITES-listed and unlisted species imported to the United States between 2009-2018 for a) proportion of the total number of imported species by threatened status, where a species is classified as: (i) threatened if their IUCN status is critically endangered (CR), endangered (EN) or vulnerable (VU); (ii) not threatened if their status is near-threatened (NT) or least-concern (LC); and (iii) unknown if their status is data-deficient (DD) or not-evaluated (NE) b) the proportion of the total number of IUCN evaluated imported species by population trend classification and c) the proportion of the total number of IUCN evaluated imported species listed as being under an ongoing threat from intentional use (code 5.1.1 intentional use: hunting & collecting terrestrial animals) for species of amphibians, birds, mammals and reptiles. Population trend and intentional use categories only compare species that have been evaluated by the IUCN (i.e., not included are those shown as Unknown in the threatened status pie chart). See Figure S1 for results of statistical comparisons.

While all species of imported birds and mammals with an IUCN Red List listing have been assessed in the past decade (as of November 2021), 40% of unlisted amphibians and 32% of unlisted reptiles had their most recent IUCN assessment completed over a decade ago (Appendix S4).

The number of new species imported to the US increased each year for all taxa during the 10-year duration of the dataset (Fig. 3). On average, new species were imported to the US at a rate of 17 per year for CITES-listed species relative to 93 for unlisted species (Fig. 3). This difference was most pronounced for reptiles where, on average, 53 new unlisted species were imported to the US per year, relative to 10 CITES-listed species (for amphibians: 14 vs. 1; for birds: 21 vs. 3; for mammals: 4 vs. 2). The number of unique species found in imports each year varied by taxa, but most taxa experienced a declining or constant rate, except for unlisted reptile species which experienced an increase in the number of unique species that appeared in imports through time (Appendix S5, Appendix S6).

**Figure 3.**
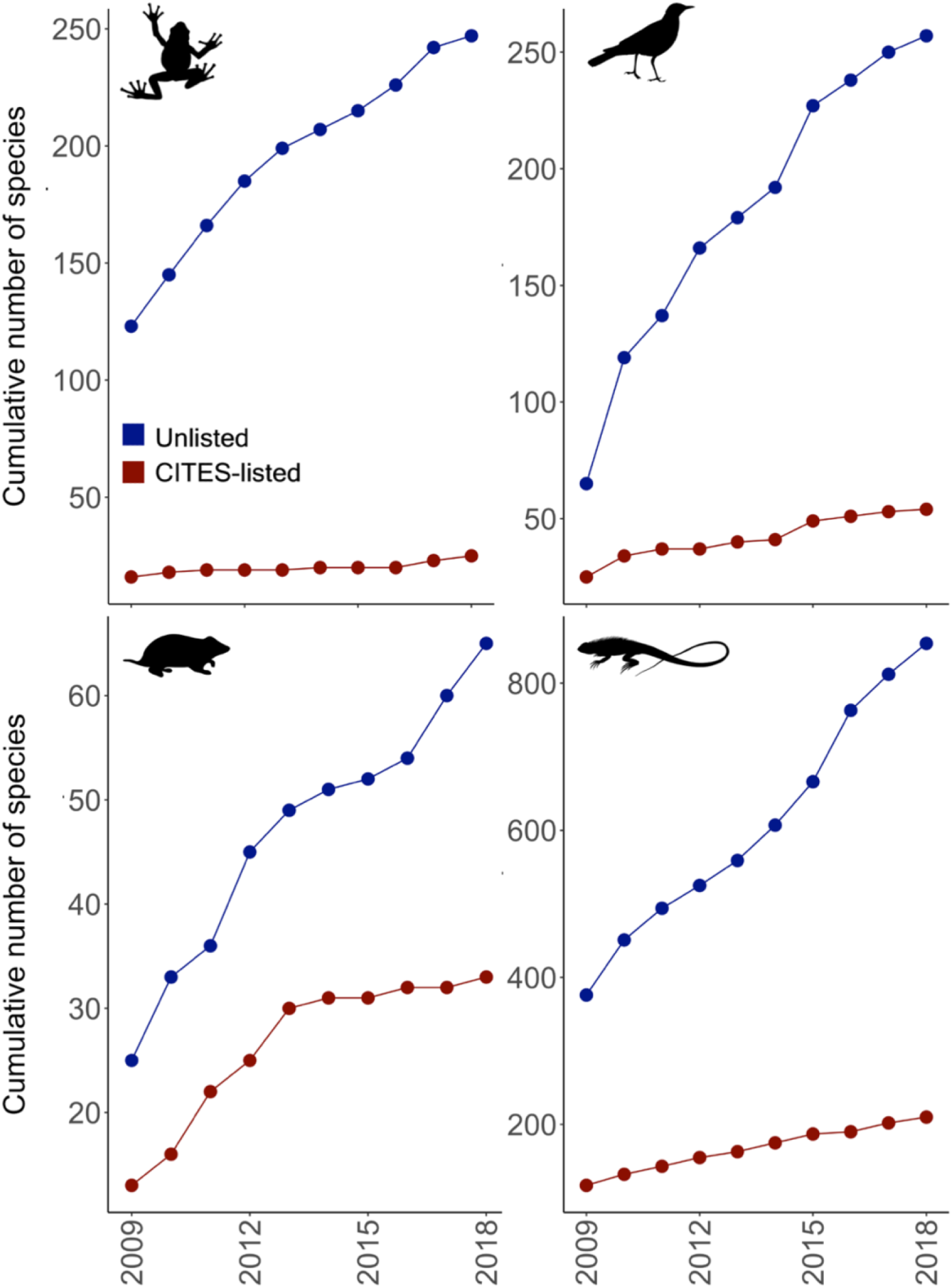
Species accumulation curves of (clockwise from top left;) amphibians, birds, reptiles and mammals imported to the United States between 2009-2018. Each point represents the number of new species imported to the US for a given year. Y-axis represents the cumulative number of species, where each species is counted for the first year it appears in US imports as either CITES-listed or unlisted.

In terms of overall import volume, eleven times as many individuals of unlisted species were imported to the US relative to CITES-listed species (8.84 million vs. 0.8 million individuals). This varied by taxa where: (i) unlisted amphibians were imported at 96 times the rate of CITES-listed species (5,378,985 versus 56,008 individuals); (ii) unlisted birds were imported at 210 times the rate (204,700 versus 973); and (iii) unlisted reptiles were imported at four times the rate (3,244,132 versus 737,785). However, when examining the distribution of the number of imports per species, only birds had significantly more per-species imports in unlisted species (Fig. 4, Appendix S8). Amphibians and mammals showed no significant difference (Fig. 4, Appendix S8). Unlisted reptiles had significantly fewer per-species imports (Fig. 4, Appendix S8). This difference largely came from imports of non-threatened reptiles, which were imported in significantly lower numbers compared with CITES-listed species, while import volumes of threatened reptiles were not significantly different (Appendix S9a, S9b Appendix S10)

**Figure 4.**
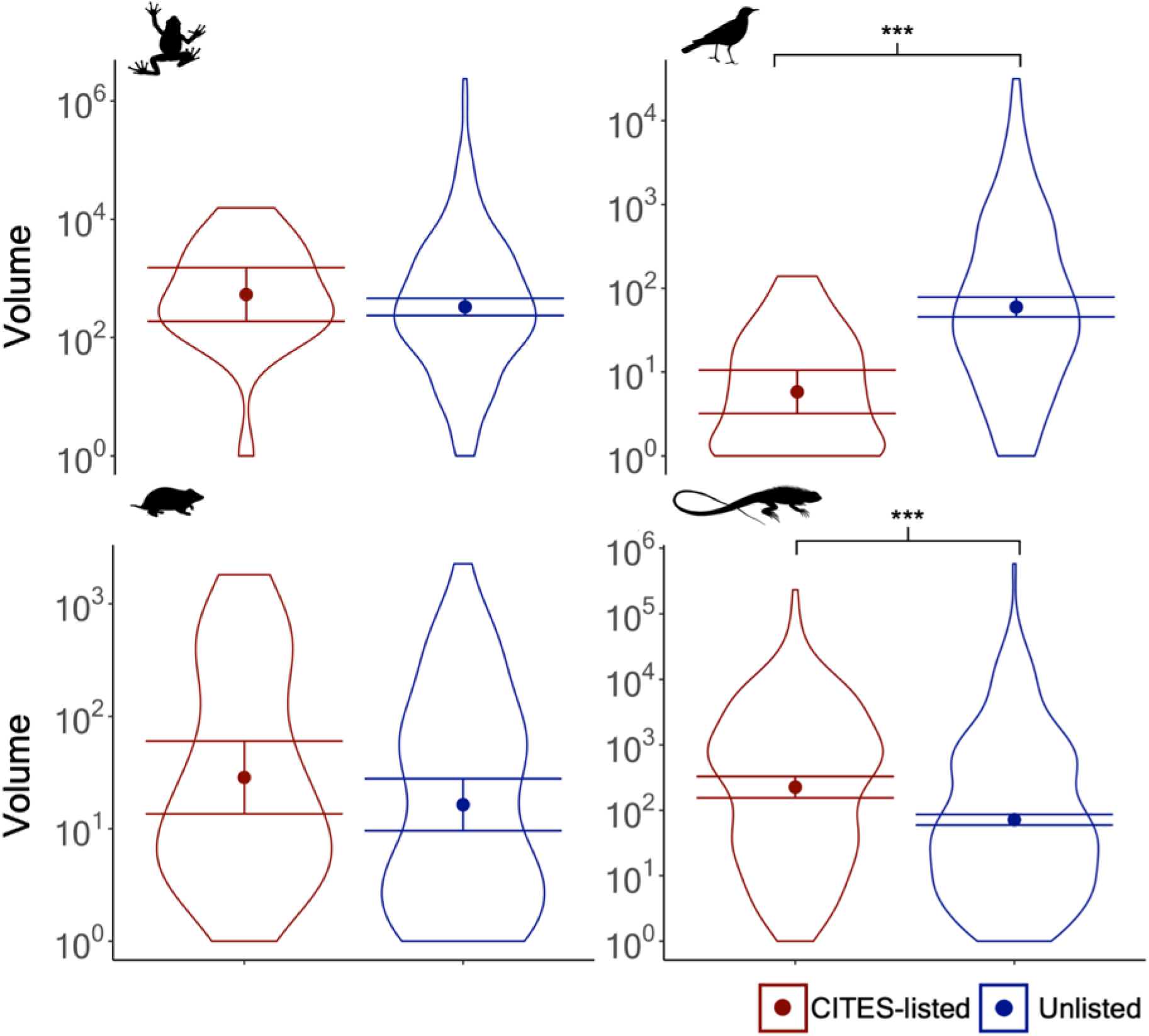
The total volume of wild-caught live imports per species, measured as individual animals for CITES-listed and unlisted species of amphibians (top left), birds (top right), mammals (bottom left), and reptiles (bottom right) entering the US between 2009-2018. Points and error bars represent the predicted median values and 95% CI’s resulting from GLM models comparing species import volumes between CITES-listed and unlisted trade. Statistically significant pairwise differences are indicated by *p*-values where *** represent *p* < 0.001. See Table S2 for further details on model results. Note the y-axis is on a log^10^ scale.

We found the countries that were top exporters of CITES-listed species were largely different from the top exporting countries of unlisted species, by both total volume and in the number of exported species (Fig. 5, Fig. 6, See Appendix 14 for complete description). For instance, amphibians had no major exporting countries in common between CITES-listed and unlisted individuals. Further, for birds, mammals, and reptiles, only a quarter of major exporting countries exported both CITES-listed and unlisted individuals (Fig. 5).

**Figure 5.**
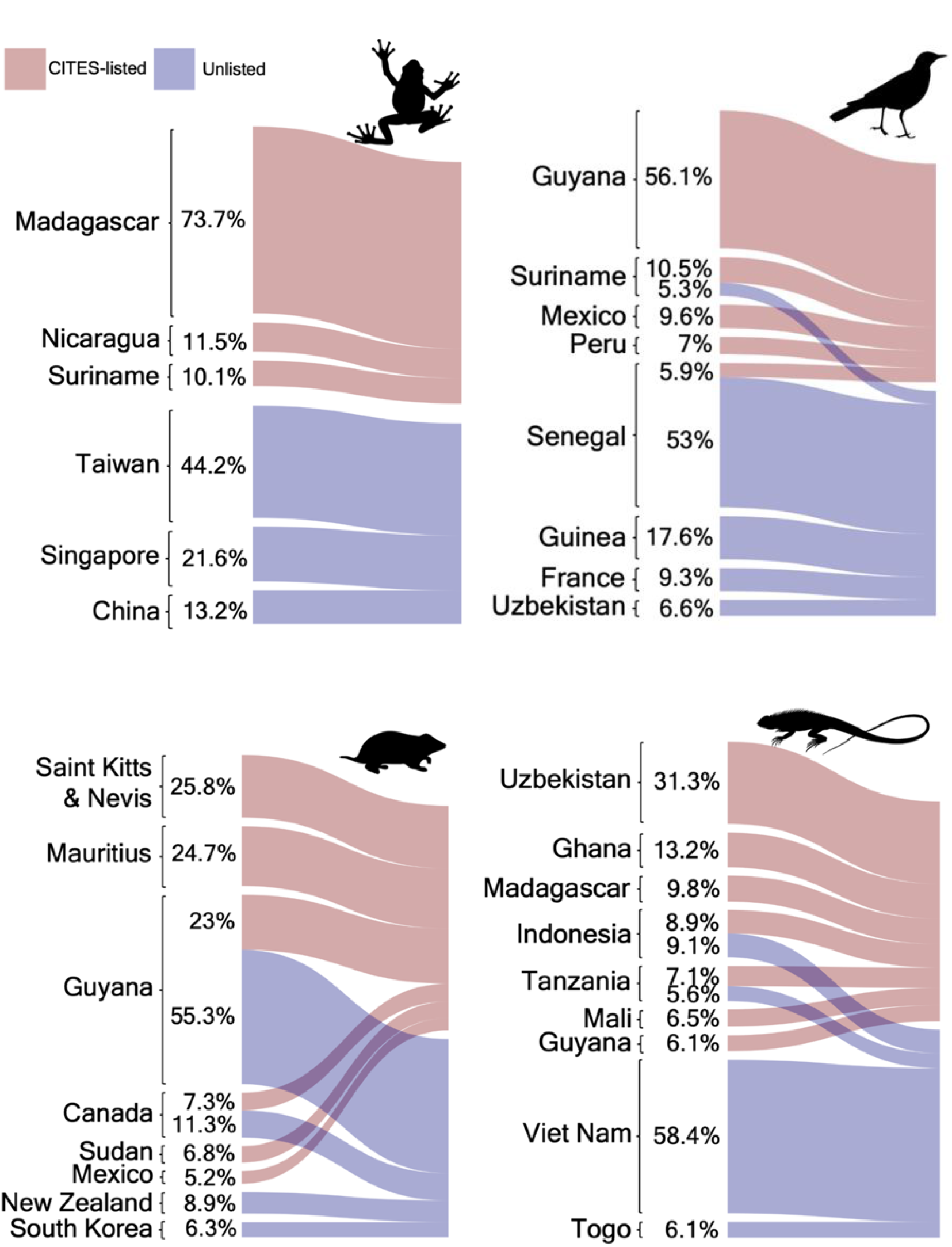
Volume of wild-caught live imports from exporting countries, measured as individual animals to the US between 2009-2018 for CITES-listed and unlisted trade. Each taxa are depicted by their silhouette: amphibians (top left), birds (top right), mammals (bottom left), and reptiles (bottom right). The width of the links corresponds to the percentage of the total trade volume each country/territory accounts for. Countries whose total trade volume ≤ 5% are not shown in the plot.

**Figure 6.**
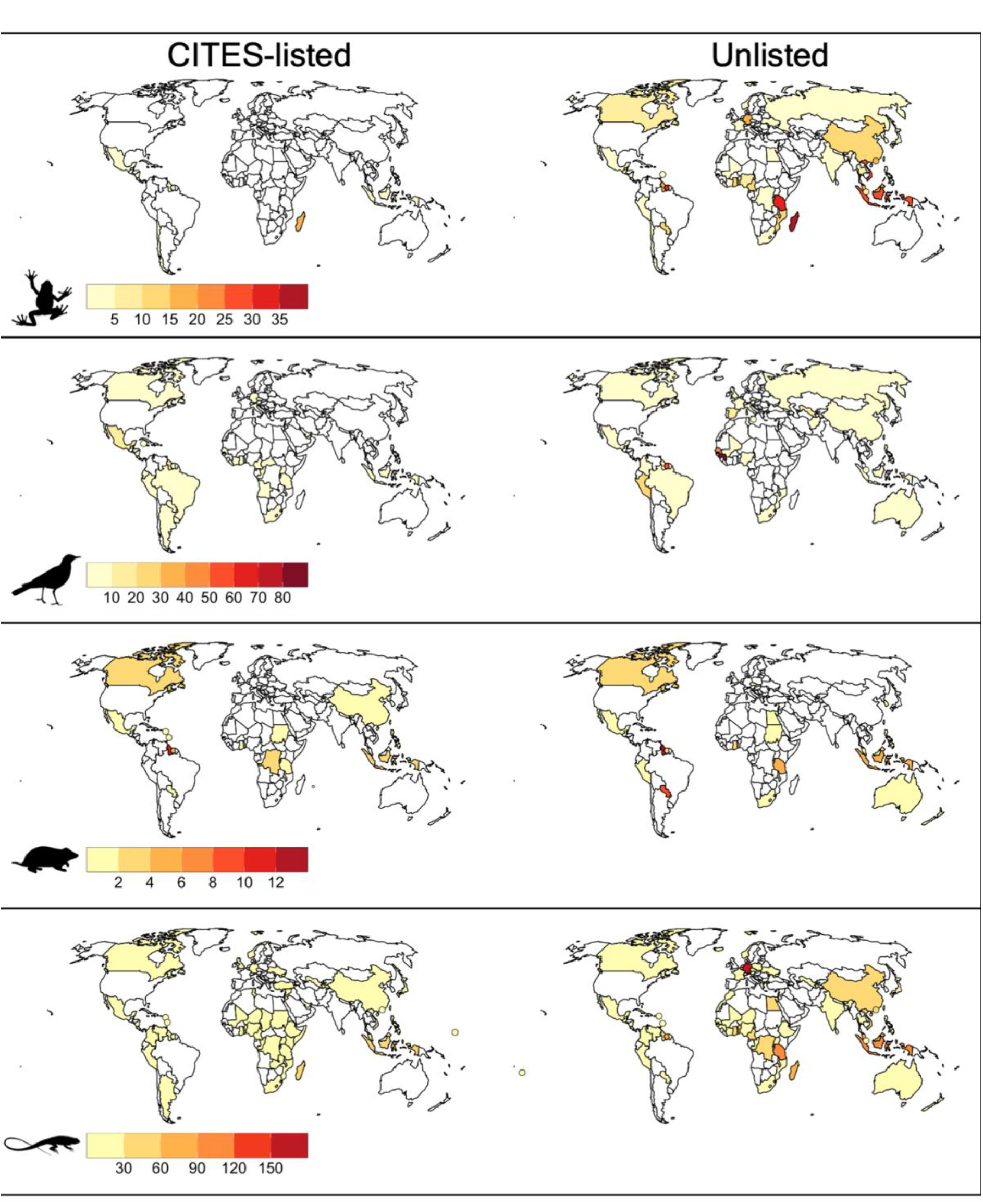
Total species richness of wild-caught live exports from each country/territory for CITES-listed (left column) and Unlisted trade (right column) entering the United States between 2009-2018 for amphibians (first row), birds (second row), mammals (third row), and reptiles (fourth row). Countries with a land mass of <5000km^2^ and a species export richness of 1 or greater are represented by points plotted at their geographic centre.

## Discussion

The global wildlife trade is a multi-billion dollar business. New species appear in trade each year, and thousands of species are traded without regulatory protections (Nellemann et al., 2014). We elucidated a thriving market for the legal trade in species not listed in the CITES appendices. This trade is increasing over time and, importantly, we identified over 350 species not protected by CITES that are facing conservation threats, yet are still being traded. We argue that the trade of species not listed in CITES demands closer attention from researchers and policymakers for both conservation and biosecurity reasons. One potential path forward is the adaptation of a global monitoring scheme to track the international trade of all wildlife.

The global trade in live terrestrial vertebrates, mainly for pets, is increasing (Bush et al., 2014; Lockwood et al., 2019), and we found a growing demand for unregulated and novel species entering the US. Importantly, the US is both the largest importer of wild-caught CITES-listed species (Liew et al., 2021) and the top importer of global wildlife trade in terms of monetary value (Andersson et al., 2021). Without the requirement for international export permits stating legal obtainment as is required with CITES species, regulating and preventing illicit and unsustainable trade in unlisted species is currently a growing concern. Evidence for ongoing international trade in unlisted reptiles and songbirds has been found in popular pet trade destinations in Asia, Europe and the US, with animals being traded in contravention to national range state protection laws (Heinrich et al., 2021a; Heinrich et al., 2021b; Janssen & Leupen, 2019; Janssen & Shepherd, 2018; Leupen et al., 2018). Recent research continues to highlight that the number of species involved in trade is greater than subsequently thought, particularly for understudied species groups popular as exotic pets (Fukushima et al., 2020; Hughes et al., 2021; Marshall et al., 2020, 2022). Of all extant described species, 17% of amphibians and 36% of reptiles have been found in trade, with unlisted species potentially being vulnerable to exploitation due to lack of trade regulations and high demand, particularly for rare and novel species, many of which have small or unknown ranges (Hughes et al., 2021; Marshall et al., 2020). Our results are consistent with these findings. Unlisted amphibians and reptiles revealed the highest import volumes and the greatest number of both species of conservation concern and species with unknown or outdated IUCN metrics. The number of novel unlisted reptile species appearing in imports averaged 53 species per year, nearly four times the average number that appeared in CITES-listed imports. Our findings suggest that the demand for amphibians and reptiles as pets is greater than ever, yet without rigorous population assessments and monitoring, we cannot be sure what level of harvest is sustainable for the majority of the species in trade (Weinbaum et al., 2013).(McRae et al., 2020; Morton et al., 2021).

For many species, the removal of individuals from the wild can have a large effect, especially for threatened species with limited ranges or small populations (Morton et al., 2021). Conservation protocols can vary significantly by species and exploitation is subject to change at any point, especially in an unchallenged situation such as unregulated trade (Eaton et al., 2015; Leupen et al., 2020). Our assessment of the sustainability of trade is limited for many species found in US imports by the uncertainty of the effects of harvest on wild populations. Many traded species are understudied, and basic data on population distribution, dynamics and threatening processes are deficient (Altherr & Lameter, 2020; Jensen et al., 2019). A third of imported unlisted species in our study showed unknown population trends (436 species, including 139 without an IUCN Red List evaluation), while 40% of unlisted amphibians and 35% of unlisted reptiles additionally had their most recent IUCN assessments completed over a decade ago, meaning data on population metrics may be potentially outdated. Outdated conservation assessments and those lacking population data may not reflect a species present circumstance, leaving the sustainability of trade in such species in doubt. Further, we found that half of unlisted species (n=652) were also not listed by the IUCN Red List as being present in international use/trade despite being present in US imports. As IUCN Red List data is frequently used in informing key international conservation management planning and policy (IUCN, 2018), the availability of accurate data on activities that may have adverse effects on wild populations, such as wild harvest for the international trade, is essential. Further assessment on the effects of trade on a by species basis, including monitoring and management of source populations, is strongly recommended to assess the sustainably of current harvesting practices and detect when potential risks emerge (Fukushima et al., 2021; McRae et al., 2020). Given the accelerating trade in unlisted species, we suggest that priority be given to expanding species-level investigation and assessment to ensure species survival is not threatened by trade.

Recent events surrounding the global SARS-CoV-2 (COVID-19) pandemic have turned the public eye towards the relationship between zoonotic disease and the wildlife trade (Andersen et al., 2020; Lu et al., 2020). Approximately 75% of the infectious diseases emerging today are zoonoses, representing a major global threat to public, agricultural, ecosystem and economic wellbeing (Gebreyes et al., 2014). The global wildlife trade provides ideal conditions for disease transmission and the emergence of novel diseases, yet there are currently no international organizations that manage the trade in wildlife based on these issues (Bell et al., 2004; Karesh et al., 2012; Shivaprakash et al., 2021). While this study did not focus on the biosecurity issues involved in the use of wildlife, the substantial volume and diversity of imports and lack of regulation implies a high risk for the transmission of emerging and re-emerging infectious diseases. For instance, a recent study involving imports of live North American bullfrog (*Rana catesbeiana)* to the US through three major national ports over five years found an overall infection prevalence of 62% in newly arrived shipments of chytridiomycosis (*Batrachochytrium dendrobatidis)*, a fungal disease infecting amphibians responsible for extensive declines in many native species worldwide (Schloegel et al., 2009). *R. catesbeiana* accounted for 44.4% (2,376,809 individuals) of total unlisted amphibian import volume during our study. In another example, the leading exporting country of unlisted wild birds in this study, Senegal, contributed 53% of the total volume of unlisted imports, had high export species richness and yet is a region that the US government department of Animal and Plant Health Inspection Service (APHIS) has recognized as affected with zoonotic diseases such as highly pathogenic avian influenza, a disease that poses a high threat to the US agricultural industry (APHIS, 2021). Exports from Senegal also included large shipments of invasive species such as the rose-ringed parakeet, (*Psittacula krameria)* (not listed in CITES and c. 19,000 individuals total exported), a species whose population establishment is a known agricultural and conservation risk in the US (Klug et al., 2019). With an average of 93 new unlisted terrestrial vertebrate species entering US imports annually, this trade deserves closer inspection purely for biosecurity concerns. Increased trade surveillance could aid in addressing current concerns regarding the need for managing wildlife trade based on zoonotic disease risks and invasion potential and further highlights the necessity for broader monitoring, data collection and regulation on a by species basis of the global wildlife trade (Sinclair et al., 2021).

No systematic alert or standard procedure exists to identify when a species may require CITES protection, and many parties have insufficient resources or incentives to implement adequate monitoring and research or to scientifically verify that harvesting is not harming wild populations (Abensperg-Traun, 2009; Challender et al., 2015). The IUCN Red List is likewise underutilized in the CITES process, with no automated action taken at CITES COPs to elect for inclusion in the appendices unlisted species that are classified by the Red List as threatened, leaving many vulnerable species unprotected (Frank et al., 2019). This oversight can affect even highly traded species, such as the Asian water dragon, (*Physignathus cocincinus),* which we found to be imported in larger total numbers than the most highly imported CITES-listed reptile species, equalling 19.3% of total unlisted reptile trade volume despite being currently listed as threatened by extinction with a severely fragmented and declining wild population trend. This species and the several hundred others we found facing similar threats should be further investigated to ascertain if trade is unsustainable or further protections are required

Various reforms have been suggested to better regulate the international wildlife trade, including ‘whitelisting’ approaches whereby only species whose trade could be assessed to be sustainable could be traded (Couzens, 2013; Macdonald et al., 2021; Marshall et al., 2020). Targeted trade bans can be an effective method at reducing disease transmission and establishment of invasive species, as well as benefitting wild populations (Cardador et al., 2019; Reino et al., 2017). In the US, the Wild Bird Conservation Act of 1992 prohibits all CITES-listed exotic birds from importation except for those included on an approved sustainable-use list (e.g., a species may be imported if wild-caught individuals were harvested following an approved management plan for sustainable use of the species). The fact that the volume and species richness of bird imports were significantly higher for unlisted trade in our study suggests that an approach that restricts imports of unlisted species with the same level of scrutiny and conservation concern as CITES-listed species could be extended to unlisted species or other at-risk groups to alleviate pressure on wild populations, while also reducing the risks that can arise from unintentional introductions of invasive species and their pathogens.

Unsustainable and at-risk trade can have severe and sustained consequences for both conservation and human livelihoods (Cardoso et al., 2021). It is therefore in the interest of the global community to ensure its future sustainability. The global scope of the challenge means significant and sustained funding, commitment, and political will would be required for an integrated approach to be successfully developed and implemented. The flow of the international trade in wildlife is predominantly from lower-income to higher-income countries (Liew et al., 2021) and therefore wealthier nations should take the lead in the funding and implementation of a universal framework to manage and report all international wildlife trade, as well as financially supporting less affluent source countries in the development of practices that are both biologically sustainable and take the livelihoods of local communities currently dependent on the global wildlife trade into account (Abensperg-Traun, 2009; Fukushima et al., 2021). Regulating the international legal trade of wildlife under a standardized electronic system would increase compliance, traceability, data accuracy and integration between countries. Such tools (e.g., eCITES; CITES Secretariat, 2017) could be used to monitor major wildlife trade hubs for early detection and prevention of over-exploitation, trading of high-risk invasives and the presence of pathogen families in species considered reservoirs for emerging zoonotic disease. A network-wide approach would be near impossible to successfully implement without systematic monitoring of all species in the international legal trade network (Sinclair et al., 2021). This is illustrated in the US importations of unlisted species. The little intersection between the major exporting countries regarding unlisted and CITES-listed imports in our study exemplifies just how broadly trade routes can differ, and that by not monitoring all legal trade, many major export hubs are potentially being underrepresented.

Our study draws much-needed attention to an often-overlooked side of the international legal wildlife trade. In highlighting the details of the wild-caught unlisted trade entering the US, a picture begins to emerge on the pervasiveness of the legal international trade in species not protected by CITES. Since CITES has no mandate for disease surveillance, humanity needs a system to monitor all legal international trade to prevent future pandemics (both human and wildlife) and avoid the massive economic costs of future invasive species (Can et al., 2019). Priority must be given to programs that promote and measure the sustainable use of wild populations on a species-by-species basis to curb trade related conservation risks (Bennett et al., 2021) and ensure that proposed regulations be grounded by robust scientific underpinning (Fukushima et al., 2021). Affluent countries where demand originates, such as the US, need to accept their role in the building of sustainable trade practices (Liew et al., 2021), including providing support for supply countries and pushing for a unified data management framework through applicable enforcement agencies to protect both humans and natural ecosystems

## Acknowledgements

We acknowledge that the land on which we conducted our research is the traditional land of the Kaurna people of the Adelaide Plains. We pay our respects to Kaurna elders past, present and emerging. We are grateful to the U.S. Fish & Wildlife Service for recording and facilitating access to the data used in our analysis. This research was partly supported by the Centre for Invasive Species Solutions (Project P01-I-002) and an Australian Research Council Discovery grant (DP210103050) to PC. FW was supported by an Adelaide University Postgraduate Research funding and stipend.

