## Supporting Information for "The demand for wildlife not protected by the CITES multilateral treaty"

Appendix S1. Mosaic plots of the deviation in conditional independence between the distribution of the percentage of CITES-listed (CL) and unlisted (UL) species imported as wild-caught live specimens classified as a) threatened (Yes (Y), No (N), Not-evaluated/Data Deficient (NE)) b) having a declining population trend (Yes (Y), No (N), Unknown (UNK)) and c) threatened by intentional use (Yes (Y), No (N)) following IUCN assessment categories for amphibians, birds, mammals and reptiles. Population trend and intentional use categories only compare species that have been evaluated by the IUCN (i.e., not included are those in the Not evaluated/Data Deficient category). Sizes of tiles in each plot are proportional to the observed frequency (number of species) for each category, which are displayed as numbers in each tile. The colour of the tiles indicates the degree relationship among the categories and responses based on Pearson residuals (Zeileis et al., 2007) where blue shading represents higher frequency than expected given independence and red shading represents lower frequency than expected by chance. *P*-values (* <0.05; ** < 0.01; *** <0.001) indicate overall Statistically significant associations between the variables and are calculated using Fisher's exact tests.

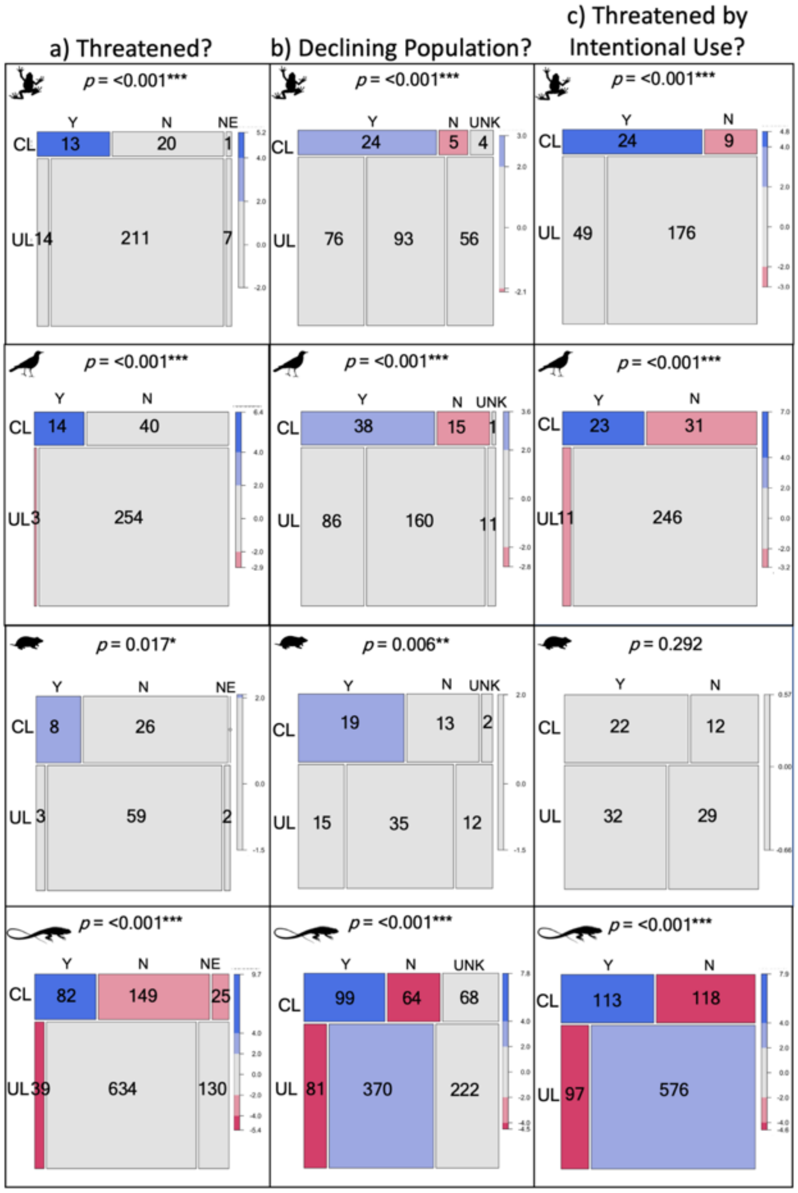
a) Threatened?
b) Declining Population?
c) Threatened by Intentional Use?
p = <0.001***
Y N NE
CL 13 20 1
UL 14 211 7
p = <0.001***
Y N UNK
CL 24 5 4
UL 76 93 56
p = <0.001***
Y N
CL 24 9
UL 49 176
p = <0.001***
Y N
CL 14 40
UL 3 254
p = <0.001***
Y N UNK
CL 38 15 1
UL 86 160 11
p = <0.001***
Y N
CL 23 31
UL 11 246
p = 0.017*
Y N NE
CL 8 26
UL 3 59 2
p = 0.006**
Y N UNK
CL 19 13 2
UL 15 35 12
p = 0.292
Y N
CL 22 12
UL 32 29
p = <0.001***
Y N NE
CL 82 149 25
UL 39 634 130
p = <0.001***
Y N UNK
CL 99 64 68
UL 81 370 222
p = <0.001***
Y N
CL 113 118
UL 97 576

Appendix S2. The overlap of the total numbers of species that are i) classified as threatened with extinction by the IUCN ii) IUCN population trend is evaluated as declining or unknown and iii) listed as being under an “ongoing” threat from wildlife trade (IUCN code 5.1.1 Intentional use: Hunting & Collecting Terrestrial Animals) that are present in wild-caught live imports to the US between 2009-2018. The total number of imported species that fall into the above listed overlapping categories compared to the total number imported are as follows and represents the order of plotting from right to left, top to bottom ; amphibians: 31/34 CITES-listed, 151/232 unlisted. Birds: 41/54 CITES-listed, 100/257unlisted. Mammals: 29/34 CITES-listed, 47/64 unlisted. Reptiles: 216/256 CITES-listed, 475/803 unlisted.

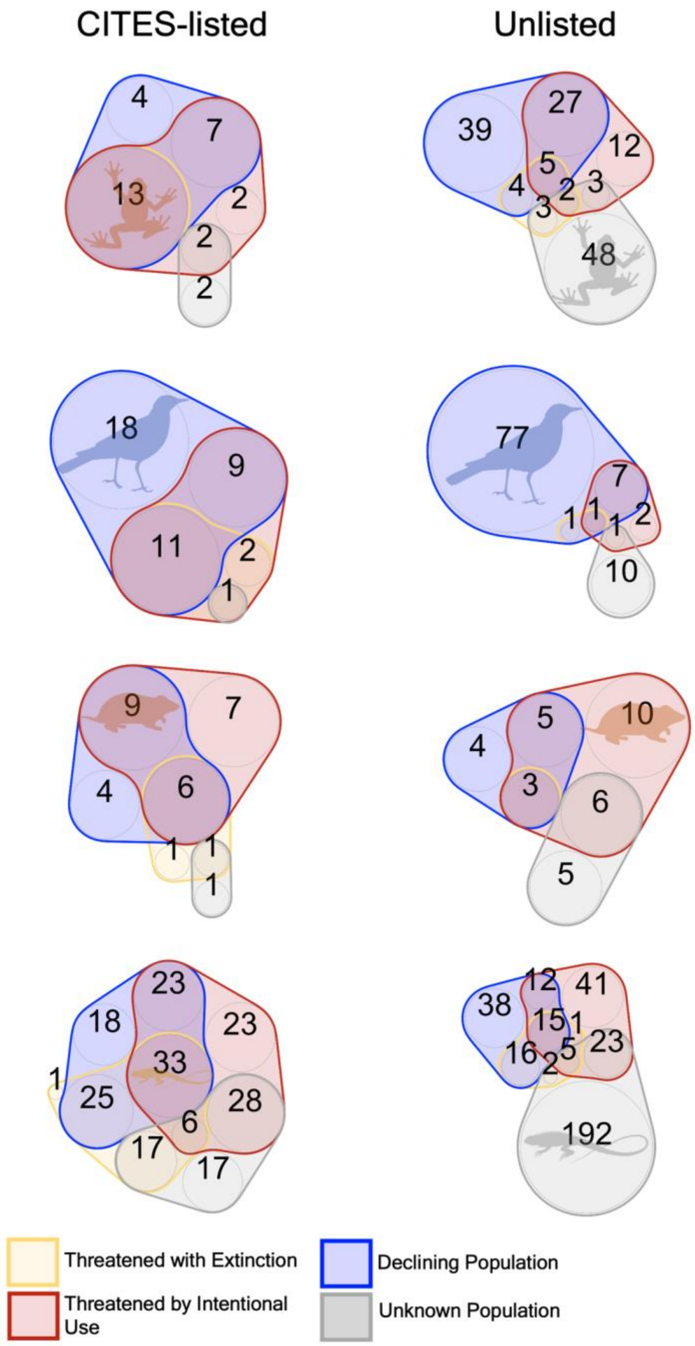

Appendix S3. The total number of IUCN evaluated species divided by trade and use status, which compares the overlap between species responses to the two IUCN categories "threatened by intentional use" and "presence in use and trade at the international level" for species of amphibians, birds, mammals and reptiles. Presence in use and trade is a classification scheme used by the IUCN to record what level of trade occurs for the species, with presence in international level trade appearing in each species listing as a binary response. Note that many species are not classified as being not present in international trade despite our records indicating they are imported to the US.

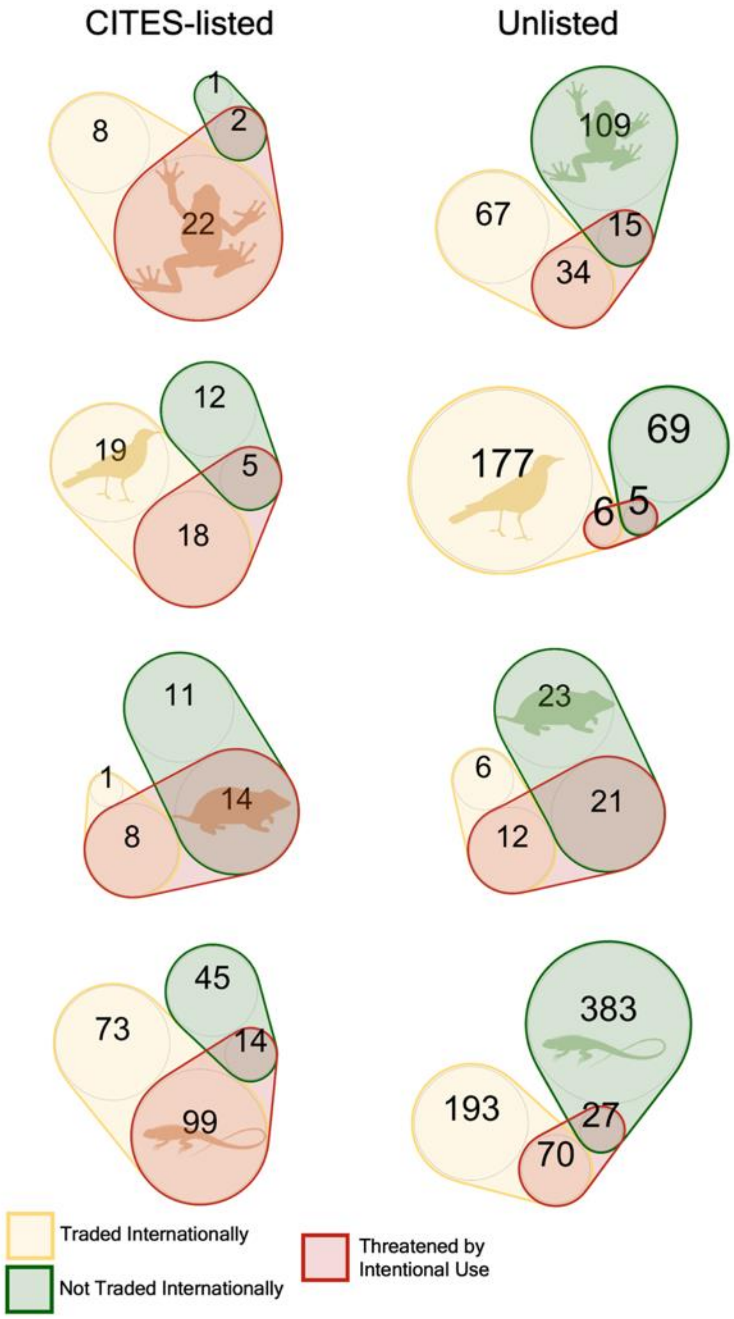

Appendix S4. Time since the most recent IUCN red list assessment was completed for CITES-listed and unlisted species imported to the United States between 2009-2018, according to their current Red List classifications (as of December 2021; (IUCN, 2021)). Bar graphs represent the number of years since the last assessment. Red list assessments are classified as out of date/needing re-assessment after ten years.

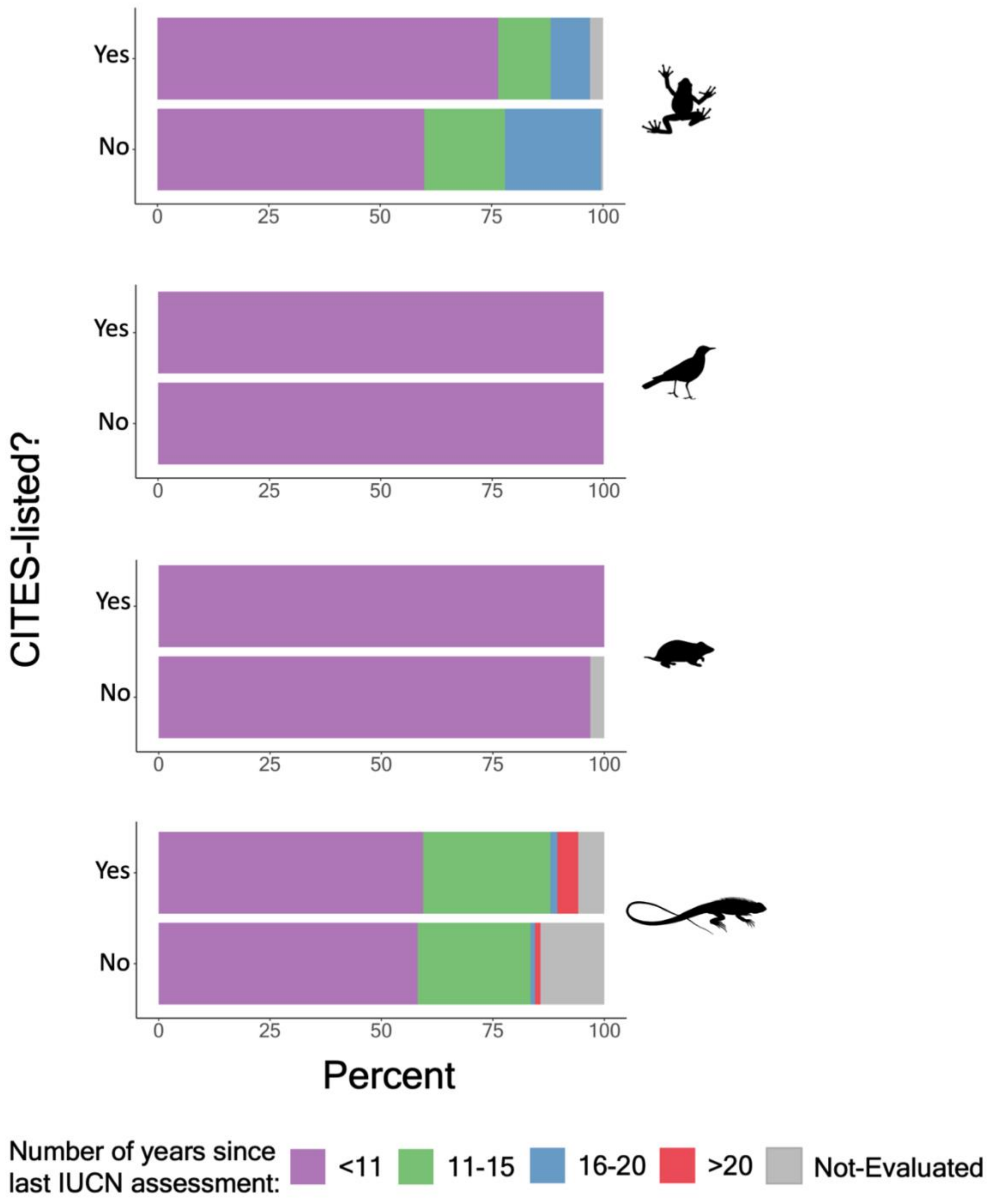

Appendix S5. The number of unique species found in CITES-listed and unlisted wild-caught live US imports by year for a) amphibians b) birds c) mammals and d) reptiles. Solid and dotted lines represent the predicted values and 95% confidence intervals of the relationship between the number of species imported through time resulting from GLM's with *p*-values equaling a statistically significant change through time (* <0.05; ** < 0.01; *** <0.001). Further model results can be located in Table S1.

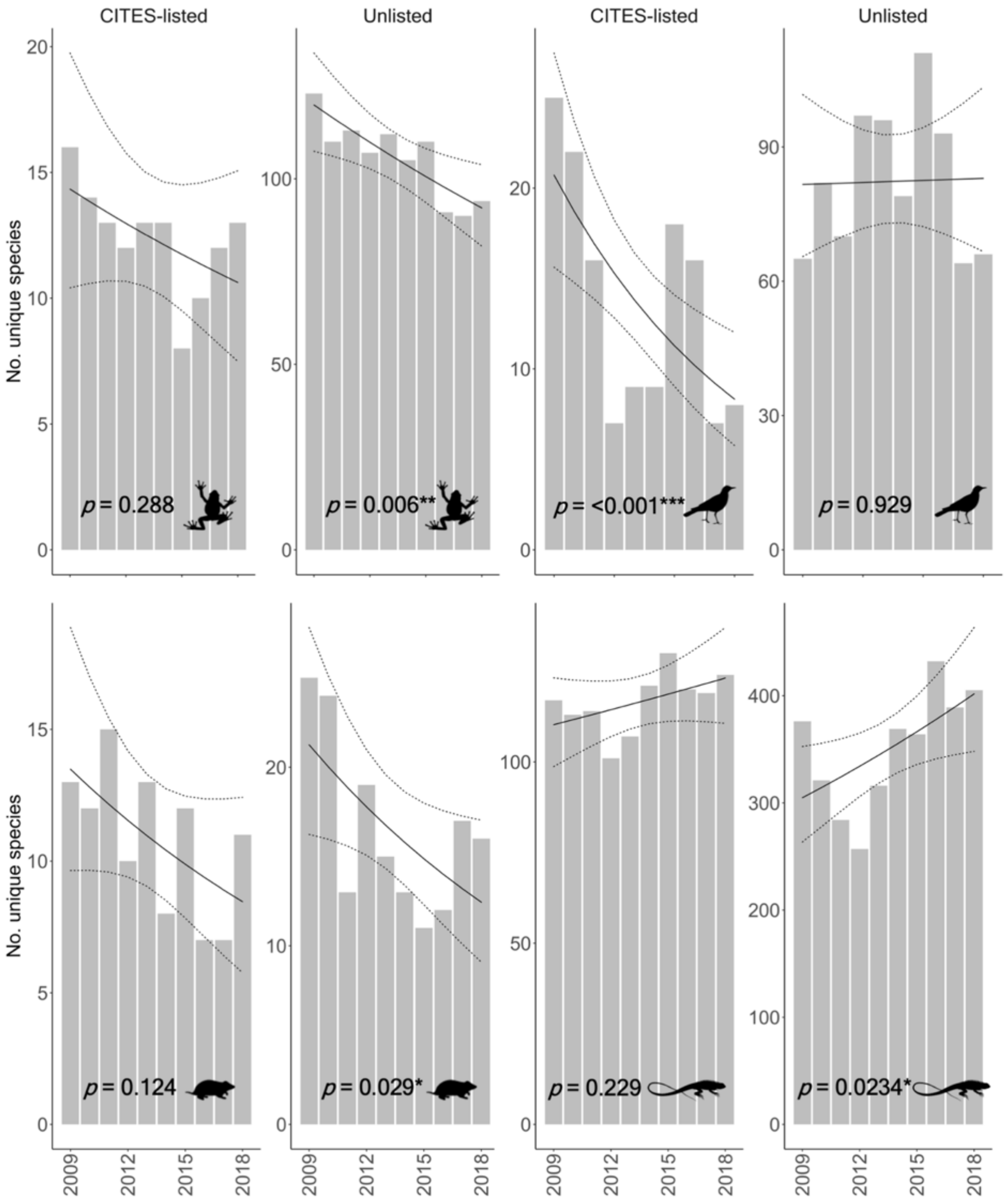

Appendix S6. Results of GLM models on the number of unique species appearing in wild-caught live CITES-listed and unlisted US imports over the 10-year sampling period (2009-2018).

| Class | CITES-listed (Yes/No) | Estimate | Std. error | z value | *p* |
|---|---|---|---|---|---|
| Amphibians | yes | -0.03 | 0.03 | -1.06 | 0.288 |
| | no | -0.03 | 0.01 | -2.73 | 0.006** |
| Birds | yes | -0.10 | 0.03 | -3.32 | <0.001*** |
| | no | 0.001838 | 0.02 | 0.09 | 0.929 |
| Mammals | yes | -0.05 | 0.03 | -1.54 | 0.124 |
| | no | -0.06 | 0.03 | -2.18 | 0.029* |
| Reptiles | yes | 0.01 | 0.01 | 1.20 | 0.229 |
| | no | 0.03 | 0.01 | 2.27 | 0.0234* |

Appendix S7. Results of the Akaike information criterion (AIC) analysis to determine the best fitting models to test for significant differences between the total volumes of all wild-caught live species and the CITES status (CITES-listed vs unlisted) of US imports across all imported species for the years encompassing 2009-2018. The models with the lowest AIC values represent the best performing models.

| Class | Model distribution | Model parameters | AIC |
|---|---|---|---|
| Amphibians | Poisson | volume ~ CITES status | 37324885.8 |
| | Negative-binomial | volume ~ CITES status | 4726.6 |
| | Gaussian | $\log^{10}$(volume) ~ CITES status | 846.4 |
| Birds | Poisson | volume ~ CITES status | 842183.7 |
| | Negative-binomial | volume ~ CITES status | 3836.6 |
| | Gaussian | $\log^{10}$(volume) ~ CITES status | 853.5 |
| Mammals | Poisson | volume ~ CITES status | 39914.0 |
| | Negative-binomial | volume ~ CITES status | 1056.5 |
| | Gaussian | $\log^{10}$(volume) ~ CITES status | 268.1 |
| Reptiles | Poisson | volume ~ CITES status | 24195742.6 |
| | Negative-binomial | volume ~ CITES status | 15541.4 |
| | Gaussian | $\log^{10}$(volume) ~ CITES status | 3375.6 |

Appendix S8. Results of the gaussian GLM models used to test for significant differences between the volume of wild-caught live CITES-listed and unlisted US imports across all imported species for the years encompassing 2009-2018, negative estimate values are associated with higher volumes of unlisted imports

| Class | Estimate | SE | t-value | *p* |
|---|---|---|---|---|
| Amphibians | 0.2 | 0.2 | 0.9 | *0.38* |
| Birds | -1.0 | 0.1 | -7.2 | <0.001*** |
| Mammals | 0.2 | 0.2 | 1.2 | 0.225 |
| Reptiles | 0.5 | 0.1 | 5.5 | <0.001*** |

Appendix S9. The total volume of wild-caught live imports distributed across imported species, measured as individual animals for CITES-listed and unlisted species that are a) i) classified as threatened by the IUCN ii) IUCN population trend is evaluated as declining and iii) listed as being under an "ongoing" threat from wildlife trade (IUCN code 5.1.1 Intentional use: Hunting & Collecting Terrestrial Animals) b) i)classified as not-threatened by the IUCN ii) IUCN population trend is evaluated as stable/increasing and c) not listed as being under an "ongoing" threat from wildlife trade, and c) i) not evaluated by the IUCN ii) IUCN population trend unknown, for species of amphibians, birds, mammals and reptiles entering the US between 2009-2018 and plotted on the log[10] scale as violin plots. Points and error bars represent the predicted values and 95% CI's resulting from GLM models comparing species import volumes between CITES-listed and unlisted trade in each category. Statistically significant pairwise differences are indicated by *P*-values (* <0.05; ** < 0.01; *** <0.001). See Table S3 for further details on model results.

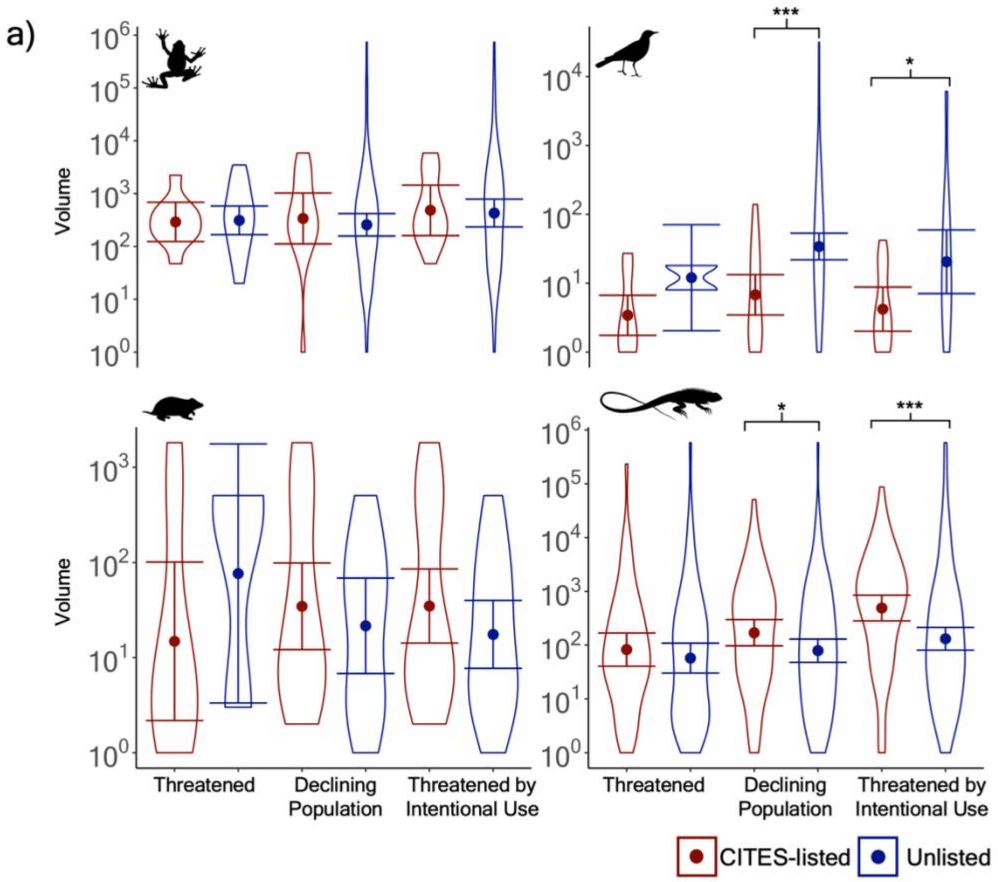

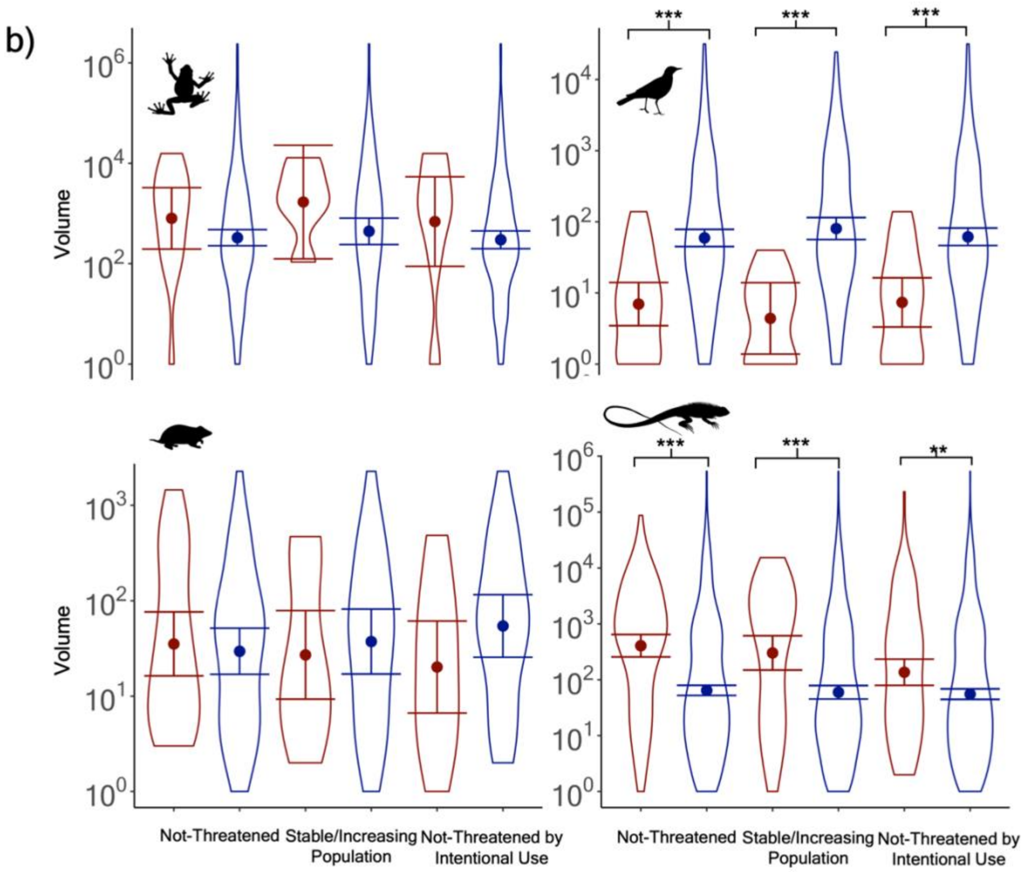
b)
Volume
Volume
***
***
***
**
Not-Threatened
Stable/Increasing Population
Not-Threatened by Intentional Use
Not-Threatened
Stable/Increasing Population
Not-Threatened by Intentional Use

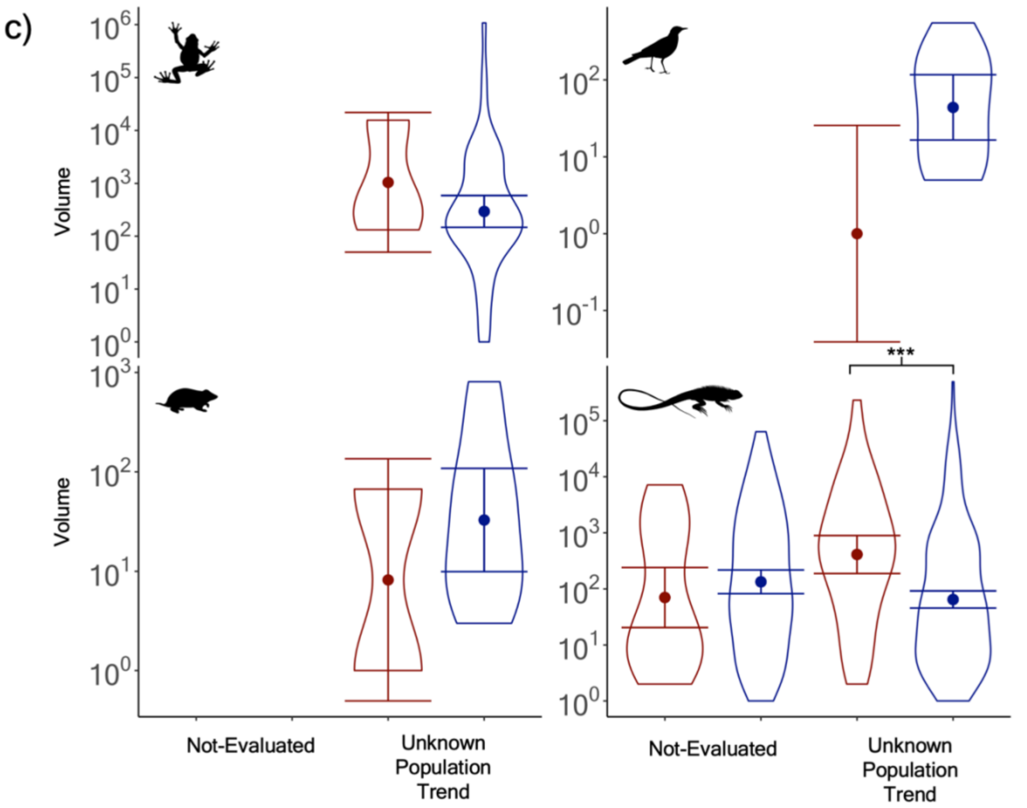
c)
Volume
Volume
***
Not-Evaluated
Unknown Population Trend
Not-Evaluated
Unknown Population Trend

Appendix S10. Results of the gaussian GLM models used to test for significant differences between the volume of wild-caught live CITES-listed and unlisted US imports across imported species for the years encompassing 2009-2018 for i) the number of imported species by threatened status, where a species is classified as threatened if their IUCN status is critically endangered (CR), endangered (EN) or vulnerable (VU), not threatened if their status is near-threatened (NT) or least-concern (LC) and unknown if their status is data-deficient (DD) or not-evaluated (NE) b) the number of IUCN evaluated imported species by population trend classification and c) the number of IUCN evaluated imported species listed as being under an "ongoing" threat from wildlife trade (code 5.1.1 Intentional use: Hunting & Collecting Terrestrial Animals).

| Class | Threatened? | Estimate | SE | t-value | *p* |
|---|---|---|---|---|---|
| | yes | -0.03 | 0.23 | -0.12 | 0.902 |
| Amphibians | no | 0.39 | 0.32 | 1.22 | 0.224 |
| | not evaluated | - | - | - | - |
| | yes | -0.55 | 0.41 | -1.33 | 0.206 |
| Birds | no | -0.93 | 0.16 | -5.70 | <0.001*** |
| | not evaluated | - | - | - | - |
| | yes | -0.71 | 0.80 | -0.89 | 0.395 |
| Mammals | no | 0.08 | 0.21 | 0.37 | 0.711 |
| | not evaluated | - | - | - | - |
| | yes | 0.16 | 0.21 | 0.77 | 0.445 |
| Reptiles | no | 0.80 | 0.11 | 7.29 | <0.001*** |
| | not evaluated | -0.28 | 0.29 | -0.97 | 0.332 |

| Class | Population Trend | Estimate | SE | t-value | *p* |
|---|---|---|---|---|---|
| | declining | 0.12 | 0.26 | 0.45 | 0.651 |
| Amphibians | stable/increasing | 0.58 | 0.58 | 1.01 | 0.317 |
| | unknown | 0.55 | 0.68 | 0.81 | 0.42 |
| | declining | -0.70 | 0.18 | -3.99 | <0.001*** |
| Birds | stable/increasing | -1.26 | 0.26 | -4.82 | <0.001*** |
| | unknown | -1.64 | 0.74 | -2.23 | 0.05 |

| | | | | | |
|---|---|---|---|---|---|
| Mammals | declining | 0.20 | 0.34 | 0.60 | 0.551 |
| | stable/increasing | -0.14 | 0.29 | -0.49 | 0.631 |
| | unknown | -0.60 | 0.66 | -0.91 | 0.382 |
| Reptiles | declining | 0.33 | 0.16 | 2.04 | 0.0431* |
| | stable/increasing | 0.70 | 0.16 | 4.27 | <0.001*** |
| | Unknown | 0.80 | 0.19 | 4.32 | <0.001*** |

| Class | Threatened by Intentional Use? | Estimate | SE | t-value | *p* |
|---|---|---|---|---|---|
| Amphibians | yes | 0.05 | 0.27 | 0.20 | 0.844 |
| | no | 0.36 | 0.46 | 0.79 | 0.429 |
| Birds | yes | -0.69 | 0.28 | -2.45 | 0.02* |
| | no | -0.92 | 0.18 | -5.03 | <0.001*** |
| Mammals | yes | 0.30 | 0.26 | 1.13 | 0.265 |
| | no | -0.43 | 0.29 | -1.48 | 0.148 |
| Reptiles | yes | 0.57 | 0.16 | 3.58 | <0.001*** |
| | no | 0.39 | 0.13 | 3.09 | 0.002** |

Appendix S11. Results of the Akaike information criterion (AIC) analysis to determine the best fitting models to test for significant differences between the total volumes of all wild-caught live species and the CITES status (CITES-listed vs unlisted) of US imports across all imported species for the years encompassing 2009-2018 by IUCN assessment category. The models with the lowest AIC values represent the best performing models.

| Class | Threatened? | Model distribution | Model parameters | AIC |
|---|---|---|---|---|
| Amphibians | yes | Poisson | volume ~ CITES status | 23710.3 |
| | | Negative-binomial | volume ~ CITES status | 437.9 |
| | | Gaussian | $log^{10}$(volume) ~ CITES status | 55.4 |
| | no | Poisson | volume ~ CITES status | 36228698.8 |
| | | Negative-binomial | volume ~ CITES status | 4089.2 |
| | | Gaussian | $log^{10}$ (volume) ~ CITES status | 747.2 |
| | not-evaluated | Poisson | volume ~ CITES status | - |
| | | Negative-binomial | volume ~ CITES status | - |
| | | Gaussian | $log^{10}$ (volume) ~ CITES status | - |
| Birds | yes | Poisson | volume ~ CITES status | 197.1 |
| | | Negative-binomial | volume ~ CITES status | 106.6 |
| | | Gaussian | $log^{10}$ (volume) ~ CITES status | 29.8 |
| | no | Poisson | volume ~ CITES status | 837646.1 |
| | | Negative-binomial | volume ~ CITES status | 3701.6 |

| | | | | |
|---|---|---|---|---|
| | | Gaussian | $log^{10}$ (volume) ~ CITES status | 814.6 |
| | not-evaluated | Poisson | volume ~ CITES status | - |
| | | Negative-binomial | volume ~ CITES status | - |
| | | Gaussian | $log^{10}$ (volume) ~ CITES status | - |
| Mammals | yes | Poisson | volume ~ CITES status | 7485.2 |
| | | Negative-binomial | volume ~ CITES status | 130.7 |
| | | Gaussian | $log^{10}$ (volume) ~ CITES status | 38.6 |
| | no | Poisson | volume ~ CITES status | 28334.7 |
| | | Negative-binomial | volume ~ CITES status | 834.5 |
| | | Gaussian | $log^{10}$ (volume) ~ CITES status | 185.3 |
| | not-Evaluated | Poisson | volume ~ CITES status | - |
| | | Negative-binomial | volume ~ CITES status | - |
| | | Gaussian | $log^{10}$ (volume) ~ CITES status | - |
| Reptiles | yes | Poisson | volume ~ CITES status | 6499340.3 |
| | | Negative-binomial | volume ~ CITES status | 1771.9 |
| | | Gaussian | $log^{10}$ (volume) ~ CITES status | 386.2 |
| | no | Poisson | volume ~ CITES status | 15751567.1 |
| | | Negative-binomial | volume ~ CITES status | 11431.9 |

| | | | | |
|---|---|---|---|---|
| | | Gaussian | $log^{10}$ (volume) ~ CITES status | 2466.2 |
| | not-evaluated | Poisson | volume ~ CITES status | 1377594 |
| | | Negative-binomial | volume ~ CITES status | 2315.377 |
| | | Gaussian | $log^{10}$ (volume) ~ CITES status | 507.7 |

| Class | Population Trend | Model distribution | Model parameters | AIC |
|---|---|---|---|---|
| Amphibians | declining | Poisson | volume ~ CITES status | 6768044.9 |
| | | Negative-binomial | volume ~ CITES status | 1736.4 |
| | | Gaussian | $log^{10}$ (volume) ~ CITES status | 297.0 |
| | stable/increasing | Poisson | volume ~ CITES status | 20720528.1 |
| | | Negative-binomial | volume ~ CITES status | 1776.0 |
| | | Gaussian | $log^{10}$ (volume) ~ CITES status | 328.2 |
| | unknown | Poisson | volume ~ CITES status | 8621953.6 |
| | | Negative-binomial | volume ~ CITES status | 1059.0 |
| | | Gaussian | $log^{10}$ (volume) ~ CITES status | 193.6 |
| Birds | Declining | Poisson | volume ~ CITES status | 374110.8 |
| | | Negative-binomial | volume ~ CITES status | 1389.3 |
| | | Gaussian | $log^{10}$ (volume) ~ CITES status | 330.5 |
| | Stable/Increasing | Poisson | volume ~ CITES status | 454603.9 |

| | | | | |
|---|---|---|---|---|
| | | Negative-binomial | volume ~ CITES status | 2290.6 |
| | | Gaussian | $\log^{10}$ (volume) ~ CITES status | 487.5 |
| | Unknown | Poisson | volume ~ CITES status | 1895.8 |
| | | Negative-binomial | volume ~ CITES status | 134.2 |
| | | Gaussian | $\log^{10}$ (volume) ~ CITES status | 29.5 |
| Mammals | Declining | Poisson | volume ~ CITES status | 14506.3 |
| | | Negative-binomial | volume ~ CITES status | 355.4 |
| | | Gaussian | $\log^{10}$ (volume) ~ CITES status | 88.1 |
| | Stable/Increasing | Poisson | volume ~ CITES status | 16055.1 |
| | | Negative-binomial | volume ~ CITES status | 464.5 |
| | | Gaussian | $\log^{10}$ (volume) ~ CITES status | 106.2 |
| | Unknown | Poisson | volume ~ CITES status | 3422.3 |
| | | Negative-binomial | volume ~ CITES status | 147.4 |
| | | Gaussian | $\log^{10}$(volume) ~ CITES status | 36.8 |
| Reptiles | Declining | Poisson | volume ~ CITES status | 5669885.1 |
| | | Negative-binomial | volume ~ CITES status | 2738.3 |
| | | Gaussian | $\log^{10}$ (volume) ~ CITES status | 556.9 |
| | Stable/Increasing | Poisson | volume ~ CITES status | 7617882.8 |

| | | | | |
|---|---|---|---|---|
| | | Negative-binomial | volume ~ CITES status | 6178.7 |
| | | Gaussian | $log^{10}$ (volume) ~ CITES status | 1374.7 |
| | Unknown | Poisson | volume ~ CITES status | 8747720.7 |
| | | Negative-binomial | volume ~ CITES status | 4281.184 |
| | | Gaussian | $log^{10}$(volume) ~ CITES status | 934.5 |

| Class | Threatened by Intentional Use? | Model distribution | Model parameters | AIC |
|---|---|---|---|---|
| Amphibians | Yes | Poisson | volume ~ CITES status | 6037377.6 |
| | | Negative-binomial | volume ~ CITES status | 1368.8 |
| | | Gaussian | $log^{10}$ (volume) ~ CITES status | 227.8 |
| | No | Poisson | volume ~ CITES status | 30872571.5 |
| | | Negative-binomial | volume ~ CITES status | 3202.2 |
| | | Gaussian | $log^{10}$ (volume) ~ CITES status | 591.3 |
| Birds | Yes | Poisson | volume ~ CITES status | 27382.2 |
| | | Negative-binomial | volume ~ CITES status | 291.6 |
| | | Gaussian | $log^{10}$ (volume) ~ CITES status | 82.3 |
| | No | Poisson | volume ~ CITES status | 812664.2 |
| | | Negative-binomial | volume ~ CITES status | 3525.8 |

| | | | | |
|---|---|---|---|---|
| | | Gaussian | $log^{10}$ (volume) ~ CITES status | 765.7 |
| Mammals | Yes | Poisson | volume ~ CITES status | 17531.9 |
| | | Negative-binomial | volume ~ CITES status | 508.0 |
| | | Gaussian | $log^{10}$ (volume) ~ CITES status | 124.1 |
| | No | Poisson | volume ~ CITES status | 15247.7 |
| | | Negative-binomial | volume ~ CITES status | 449.8 |
| | | Gaussian | $log^{10}$(volume) ~ CITES status | 98.1 |
| Reptiles | Yes | Poisson | volume ~ CITES status | 10011423.7 |
| | | Negative-binomial | volume ~ CITES status | 3451.3 |
| | | Gaussian | $log^{10}$ (volume) ~ CITES status | 659.0 |
| | No | Poisson | volume ~ CITES status | 10864585.5 |
| | | Negative-binomial | volume ~ CITES status | 9700.0 |
| | | Gaussian | log10(volume) ~ CITES status | 2186.3 |

Appendix S12. Exporting countries/territories whose contribution to total trade volume equals >= 5%  for amphibians, birds, mammals, and reptiles and the top two species by total export volume for each respective country (see separate excel document).

Appendix. S13 The total annual import volumes of individual animals entering the US for the top five traded species by total volume (summed across the total study period) for a) CITES-listed and b) unlisted species of amphibians, birds, mammals and reptiles.

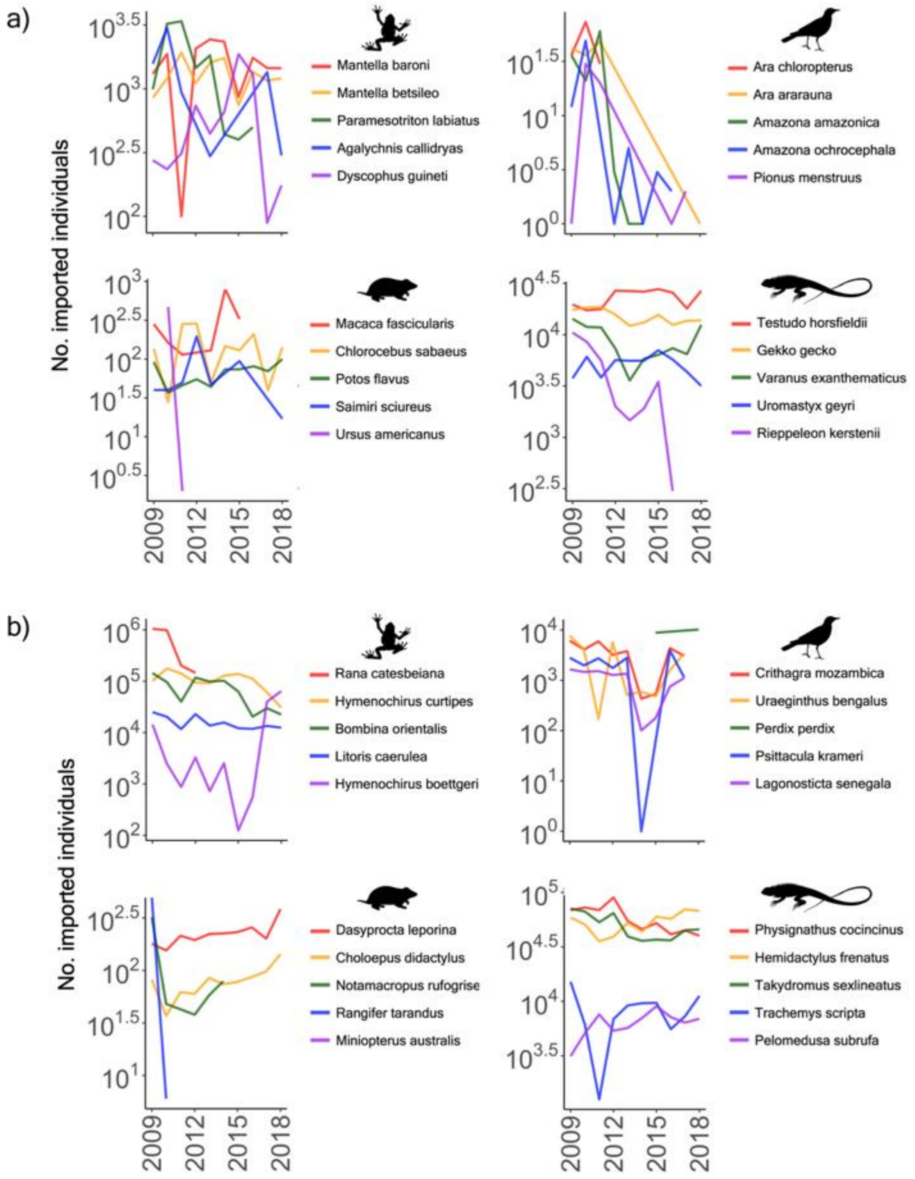

Appendix 14: Exporting Countries

For CITES-listed trade of amphibians entering the US, Madagascar was both the largest exporter by volume and exporter of the greatest number of species (Fig. 5, Fig. 6). The greatest volume of unlisted amphibians came from Taiwan and Singapore, both countries where 99% of their trade volume came from singular (*Rana catesbeiana*) or two (*Hymenochirus curtipes*, *Hymenochirus boettgeri*) species respectively (Table S4). While exporters with high species richness of exports (but low total trade volumes of <5%) included Suriname, Madagascar, Tanzania, Southeast Asia, and Germany.

Guyana was the largest exporter of CITES-listed birds by volume (56.1% of trade, Fig. 5). Guyana also exported the highest number of species, along with Mexico, which also contributed 9.6% of total trade volume (Fig. 5, Fig. 6). Species richness in unlisted bird trade was highest from Suriname and the West African countries of Guinea and Senegal, who were also the two largest contributors to total unlisted trade volume (Fig. 5, Fig. 6).

The three largest exporters of mammals by volume were Guyana, Saint Kitts and Nevis and Mauritius (Fig. 5). Guyana also exported the greatest number of species (Fig. 6) whilst the export volumes of Saint Kitts & Nevis and Mauritius was composed entirely of Chlorocebus monkeys (*Chlorocebus sabaeus* and *Chlorocebus aethiops*) and macaques (*Macaca fascicularis*) respectively (Table S4). Guyana was also the top exporter in both volume and species richness for unlisted imports, while exporter species richness was also high for the countries of South Africa and Paraguay, with both countries contributing less than 5% to total trade volume (Fig. 5, Fig. 6).

CITES-listed exporters of reptiles with both a high volume of exports and a high species richness included Indonesia and the East African countries of Madagascar and Tanzania (Fig 5, Fig 6). Uzbekistan was the top exporter by volume, but only exported one species, the Russian tortoise (*Testudo horsfieldii,* Table S4), whilst other countries with high export species richness such as Germany had relatively small export volumes (0.1%). Germany and the neighbouring Netherlands also had a high diversity of exports for unlisted species (Fig. 6), with similarly small total export volumes (<1%). Exporter species richness was also high for Tanzania, Indonesia, and Malaysia whilst the largest exporter by volume was Viet Nam, which exhibited medium level species export richness (Fig. 5, Fig. 6).
